# Lung mesenchymal cell diversity rapidly increases at birth and is profoundly altered by hyperoxia

**DOI:** 10.1101/2021.05.19.444776

**Authors:** Fabio Zanini, Xibing Che, Nina E Suresh, Carsten Knutsen, Paula Klavina, Yike Xie, Racquel Domingo-Gonzalez, Min Liu, Robert C Jones, Stephen R Quake, Cristina M. Alvira, David N. Cornfield

## Abstract

Lung mesenchymal cells play an essential role in development and at birth, as the lung moves from a fluid-filled to an oxygen-rich environment with a stable gas-liquid interface. The molecular details and cellular changes accompanying this highly coordinated process remain incompletely understood. Therefore, we performed single cell transcriptomics and in-situ imaging of the developing lung in both health and disease to characterize the spectrum of mesenchymal cell states prior to the onset of air-breathing life through late alveolarization to gain insight into their role in orchestrating tissue maturation. We found that cell type diversity in the mesenchymal compartment increases rapidly during normal development but is delayed during neonatal exposure to 80% O_2_ hyperoxia, a model for bronchopulmonary dysplasia. This study identifies the molecular transitions between populations of mesenchymal cells at discrete developmental time points across fibroblast, smooth muscle, and mural compartments and elucidates the global and cell type-specific effects of neonatal hyperoxia, including the emergence of *Acta1*+ cells which are absent in normoxic neonatal lungs. These granular insights hold the promise of targeted treatment for neonatal lung disease, which remains a major cause of infant morbidity and mortality across the world.

## Introduction

Lung development requires coordinated interactions between multiple cell types in the epithelial, endothelial, immune and mesenchymal compartments [1,2]. *In utero,* the lung is fluid filled and the pulmonary circulation develops in a relatively low oxygen tension environment with low blood flow and high pressure [3]. Within moments after birth, the perinatal lung undergoes a remarkable transition that enables gas exchange with establishment of a gas-liquid interface, a ten-fold increase in pulmonary blood flow and a marked decrease in pulmonary arterial pressure [4,5]. After birth, the distal lung undergoes structural remodeling including the formation of alveoli by secondary septation and a transition from a double to single capillary layer to increase gas exchange efficiency [6].

Lung development requires coordinated interactions between multiple cell types in the epithelial, endothelial, immune and mesenchymal compartments [1,2]. Lung mesenchymal cells (MC) play a central role in secondary septation, thinning of the interstitium, and transmitting mechanical forces that promote alveolarization [7]. Although the lung mesenchyme includes multiple, distinct cell types, knowledge surrounding the degree of cellular heterogeneity, dynamic changes in gene expression, and cell-cell interactions during alveolarization remains limited.

Insight into the lung mesenchyme during early postnatal life has significant implications for lung injury and regeneration, across the lifespan, but especially in neonates. Since the lung continues to develop in late pregnancy and through the first decade of life [8], premature infants are uniquely susceptible to life-threatening lung disorders, including bronchopulmonary dysplasia (BPD), a lung disease characterized by compromised alveolarization [9]. Given the importance of the mesenchyme in both physiologic development and developmental disease, we sought to interrogate the lung mesenchyme relative to cellular diversity, dynamic changes in gene expression and cell-cell communication in the perinatal lung, during alveolarization, and in the context of hyperoxia-induced lung injury, a preclinical model of BPD [10].

In this report, we combined single cell transcriptomics (scRNA-Seq) with fluorescent multiplexed in situ hybridization (FISH) and computational analyses to characterize changes in composition, localization, and function of MC in the murine lung from just before birth through the first three weeks of postnatal life. Moreover, we applied the same strategies to cells derived from neonatal murine lung after 7 days of hyperoxia (80% oxygen). Consistent with prior studies, MC fell into three broad categories: fibroblasts, airway smooth muscle (ASM)/myofibroblasts (MyoF), and mural cells. Overall, the present strategy identified 14 distinct subtypes, some of which restricted to perinatal developmental stages. Remarkably dynamic changes in cell phenotype occurred at birth as separate precursor cells for either fibroblasts or ASM/MyoF led to distinct but related cell subtypes between E18.5 and P1. At P7, early ASM cells were localized not only around the airways, but also in the distal lung, spatially proximate to, but transcriptionally distinct from, MyoF. At P7 lung MC were most proliferative and corresponded to the emergence of a novel proliferating population of MyoF. Hyperoxia blocked the maturation-related transcriptomic progression of MC, mirroring the structural arrest in lung development that characterizes BPD in infants, and markedly decreased pericyte and myofibroblast abundance and proliferation. Further, both single-cell and bulk transcriptomics as well as by FISH imaging identified previously undescribed *Acta1+*MC in multiple hyperoxia-exposed mice. These data demonstrate that the neonatal lung mesenchyme possesses a high degree of cellular diversity and changes dynamically with development, with hyperoxia causing both generalized transcriptomic arrest and the specific emergence of novel, previously undescribed MC.

## Results

### The perinatal lung mesenchyme is composed of three groups of cell types

Single cell RNA-Seq (scRNA-Seq) data were generated from eight mice at different stages of perinatal development, with two mice (one female and one male) from four points during development, embryonic day 18.5 (E18.5; early saccular stage), postnatal day of life 1 (P1; late saccular), P7 (early alveolar) and P21 (late alveolar). Lung tissue was resected, perfused, and dissociated as described previously [11]. Fluorescence activated cell sorting (FACS) was used to enrich mesenchymal cells by depleting cells positive for CD326, CD31, and CD45 (**Figure 1A**). Libraries were generated in 384-well plates, combining medium cell throughput with excellent per-cell capture sensitivity as demonstrated previously [11]. A total of 5,493 high-quality cells at an average depth of 980,000 read pairs per cell were analyzed (see Methods), yielding a much larger number of detected genes per cell compared to any other study of the perinatal lung (**Figure 1 Supplementary 1**).

**Figure 1.**
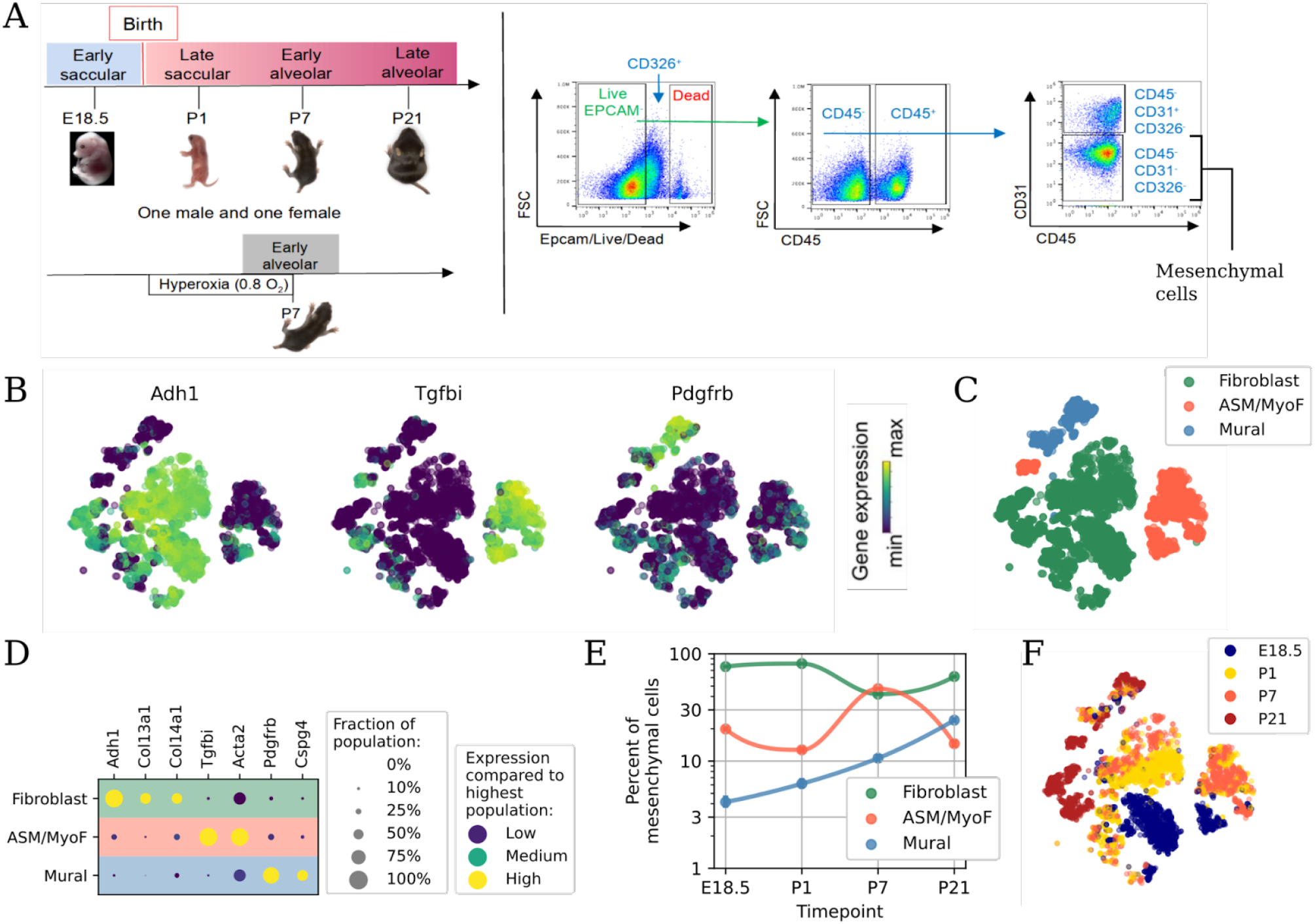
The perinatal lung mesenchyme is populated by three groups of cell types. (A) Experimental design including time points sampled, tissue dissociation, and cell sorting to isolate mesenchymal cells. (B) Embedding of lung mesenchymal cells, colored by relative expression of four distinguishing marker genes. (C) As in B, but colored by cell type group. (D) Dot plot of representative marker genes for each group. (E) Relative abundance for each group in normal development. (F) As in B, but colored by time point of each cell. At this level of resolution, airway smooth muscle (ASM) and myofibroblasts (MyoF) belong to the same group (ASM/MyoF).

Mesenchymal cells are among the most diverse but least well characterized cells of the body [12]. An initial representation of the perinatal lung mesenchyme was constructed by computing a t-distributed stochastic neighbor embedding (t-SNE)[13] and categorizing each cell based on expression of established marker genes (**Figure 1B**). Three groups of mesenchymal cell types were defined (**Figure 1C**). Fibroblasts were marked by *Adh1* [14,15]. A heterogeneous group of cells including both airway smooth muscle cells and myofibroblasts (ASM/MyoF) was marked by a shared expression of *Tgfbi* [16]. Finally, mural cells were marked by *Pdgfrb* [17] (**Figures 1D** and **Table S1**). Across four time points spanning E18.5 until P21, fibroblasts constituted the majority of cells except at P7. ASM/MyoF abundance peaked at P7, and mural cell abundance slowly increased over time (**Figure 1E**). Both cell type abundance and gene expression changed dynamically with maturation (**Figure 1F**). Remarkably, no cluster coexpressed of *Tgfbi* and *Pdgfrb* (**Figure 1D**), previously described as vascular smooth muscle cells (VSM) markers [18], while ASM/MyoF embedded in very distinct locations at perinatal times vs P21 (**Figures 1C and 1F**), arguing for the importance of identifying distinct types of smooth muscle, unlike done in some publications [19–22].

### Perinatal and adult mesenchymal cell subtypes are transcriptionally distinct

Annotation of fine-grained cell subtypes within each group was particularly challenging given the inconsistencies in the published literature [19,20,22–24]. Leiden clustering [25] resulted in 14 distinct clusters (**Figure 2A**). To annotate them, harmonized co-embedding with an adult cell atlas, Tabula Muris Senis (TMS, **Figure 2B**) [20], was combined with marker gene analysis (**Figure 2C**). Interestingly, while clusters composed of predominantly P21 cells (4, 6, 9, 12, 13, 14) were consistent with the adult atlas, clusters derived from earlier developmental time points (clusters 1-3, 5, 7-8, 10-11) embedded separately from the atlas in spite of the underlying batch-balanced k-nearest neighbor network [26], supporting the notion that perinatal lung cells possess transcriptomic profiles distinct from adult. Clusters 1-6 were consistent with fibroblasts, clusters 7-11 included ASM and myofibroblasts, and clusters 12-14 mural cells. In the co-embedding, clusters 13 and 14 co-localized with the atlas pericytes, cluster 4 with atlas alveolar fibroblasts and cluster 6 with atlas adventitial fibroblasts (**Figure 2B**).

**Figure 2.**
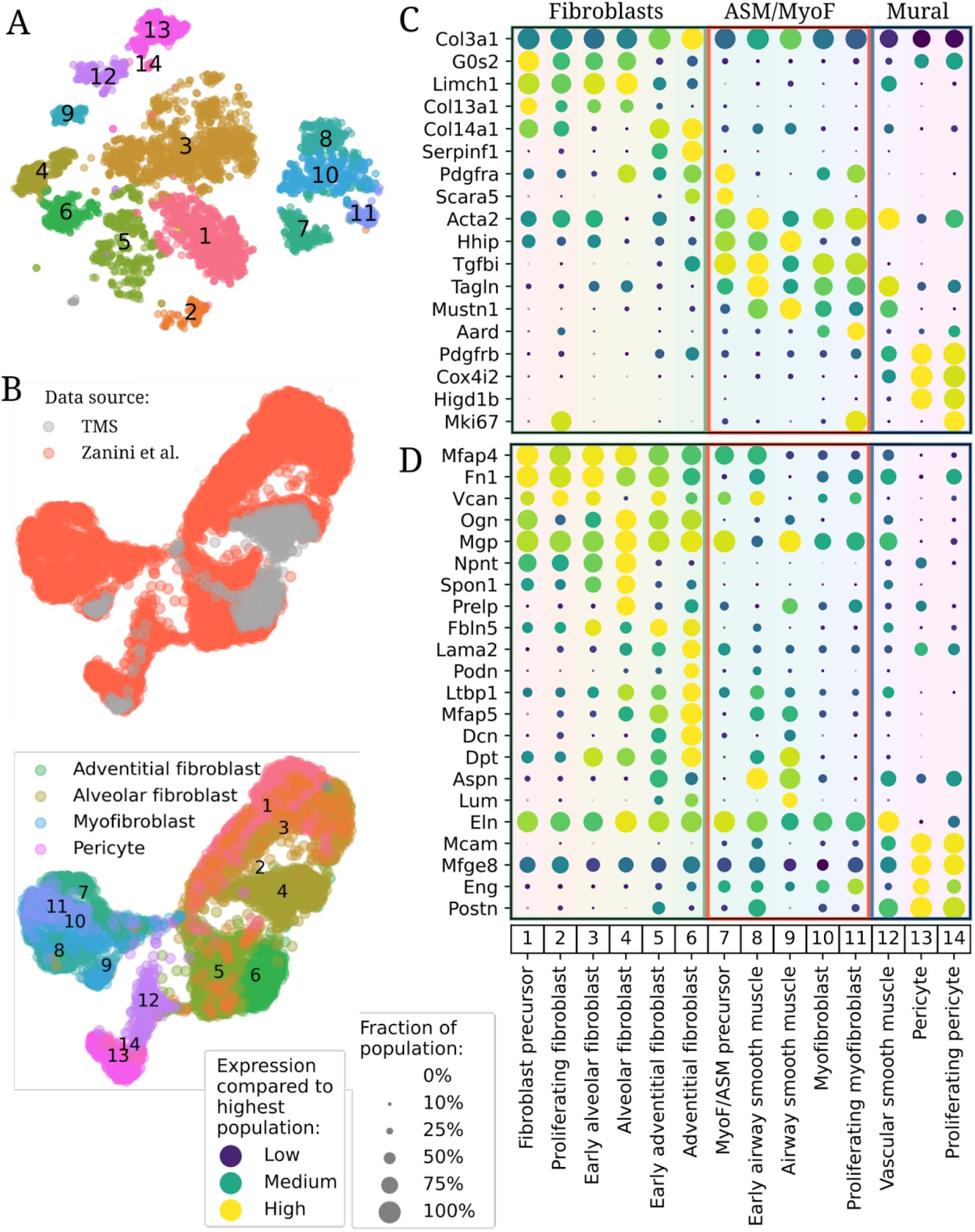
Harmonized fine grained composition of the perinatal mesenchyme. (A) Embedding as in Figure 1B, but colored by fine cell type and annotated with cluster numbers 1-14. This color code is used in subsequent figures. (B) Harmonized embedding of perinatal mesenchymal cells with mesenchymal cells from TMS colored by source (top panel, gray TMS cells drawn on top) and cell type (bottom, merging concordant cell types between our data and TMS). (C-D) Dotplots of all mesenchymal cell types in this study for (C) marker genes and (D) components of the extracellular matrix. TMS: Tabula Muris Senis.

The annotation of clusters 9 and 12 could not be annotated on the basis of the adult atlas alone. For cluster 12, which co-embedded between smooth muscle-like cells and pericytes (**Figure 2B**), the coexpression of both smooth muscle (*Tagln, Acta2)* and mural *(Prgfrb)* genes (**Figure 2C**) identified this population as vascular smooth muscle, a cell type that, while not annotated in the adult atlas, is clearly present based on lung histology [27] and in a previous single-cell transcriptomic study [28]. Cluster 9 derived from P21 mice (**Figure 1E**) and colocalized with a group of atlas myofibroblasts (**Figure 2B**). However, myofibroblasts emerge in adult mice under pathological conditions only (e.g. injury) [29] and cluster 9 expressed *Hhip*, an established ASM marker [30] but lacked *Pdgfra,* a myofibroblast marker, therefore these cells were annotated as ASM (**Figure 2C**). In turn, this led to the tentative annotation of cluster 8 *(Hhip+ Pdgfra-)* as early ASM and cluster 10 (*Hhip-Pdgfra+)* as myofibroblasts (**Figure 2C**). Both clusters comprised mostly cells from P1 and P7. At E18.5, a single *Tgfbi+* cluster was found (cluster 7). It coexpressed *Hhip* and *Pdgfra* and was therefore annotated as ASM/MyoF precursors. Clusters 2, 11, and 14 expressed *Mki67* and were therefore labeled as proliferating fibroblasts, myofibroblasts, and pericytes, respectively (**Figure 2C**). To contextualize our cell type definitions, we computed dot plots of marker genes from recent publications against our cell types [23,31] (**Figure 2 Supplementaries 1 and 2**).

To further buttress the physiologic relevance of the transcriptomic findings, expression of genes associated with extracellular matrix (ECM) production and remodeling were evaluated (**Figure 2D**). Fibroblasts shared ECM components *Mfap4, Fn1, Vcan,* and *Ogn.* In alveolar fibroblasts (AlvF), subtype-specific expression included *Spon1* which encodes a secreted adhesion protein [32]. Mature and Early Adventitial fibroblasts (AdvF) expressed *Mfap5,*which contributes to tissue elasticity [33], and *Dcn,* which confers resistance to tissue compression, while only adventitial fibroblasts from P21 mice expressed *Podn,* encoding a molecule that constrains smooth muscle cell proliferation and migration [34]. Early ASM highly expressed *Aspn,* which inhibits canonical TGFβ and SMAD signaling [35] and competes with *Dcn* for collagen binding [36], while ASM from P21 animals had reduced expression. However, mature ASM expressed *Lum,* which regulates collagen fibril organization. Interestingly, Early ASM, found in P1 and P7 mice, did not express *Lum,* suggesting other genes might regulate collagen fibrils at those time points [37,38]. Pericytes, which modulate vascular tone, endothelial cell (EC) migration and angiogenesis [39], expressed *Mcam, Mfge8, Postn,* and *Eng.* Vascular smooth muscle had the highest expression of elastin (*Eln*), while all other cell types except pericytes expressed some level of *Eln.*

### Fibroblast and ASM/MyoF precursors differentiate at the onset of air-breathing life

The cell composition of the lung changed dramatically between E18.5 and P1, a time window of just 48 hours. Fibroblast precursors, the most abundant mesenchymal type at E18.5, disappeared by P1 (**Figure 3A**), concomitant with the appearance of two distinct types of postnatal fibroblasts (**Figure 3B** and **Table S3**) distinguished by expression of either *Col13a1* or *Col14a1* (**Figure 3C**), resembling adult alveolar and adventitial fibroblasts, respectively. *Col13a1* and *Col14a1* were coexpressed in the E18.5 precursors, suggesting a common origin (**Figure 3C**), a hypothesis corroborated by trajectory analysis via diffusion maps, pseudotime, and RNA velocity (**Figure 3 Supplementary 1**). In terms of ECM-associated gene expression, fibroblast precursors appeared slightly more similar to alveolar fibroblasts (**Figure 2D** and **Figure S4**). Differentially expressed genes between embryonic and a balanced mix of postnatal fibroblasts showed a statistical enrichment of mesenchyme-, ECM-, and vascular development-associated pathways in both groups (**Figure 3D**, red/blue bars upregulated in precursors/postnatal cells, respectively) [40].

**Figure 3.**
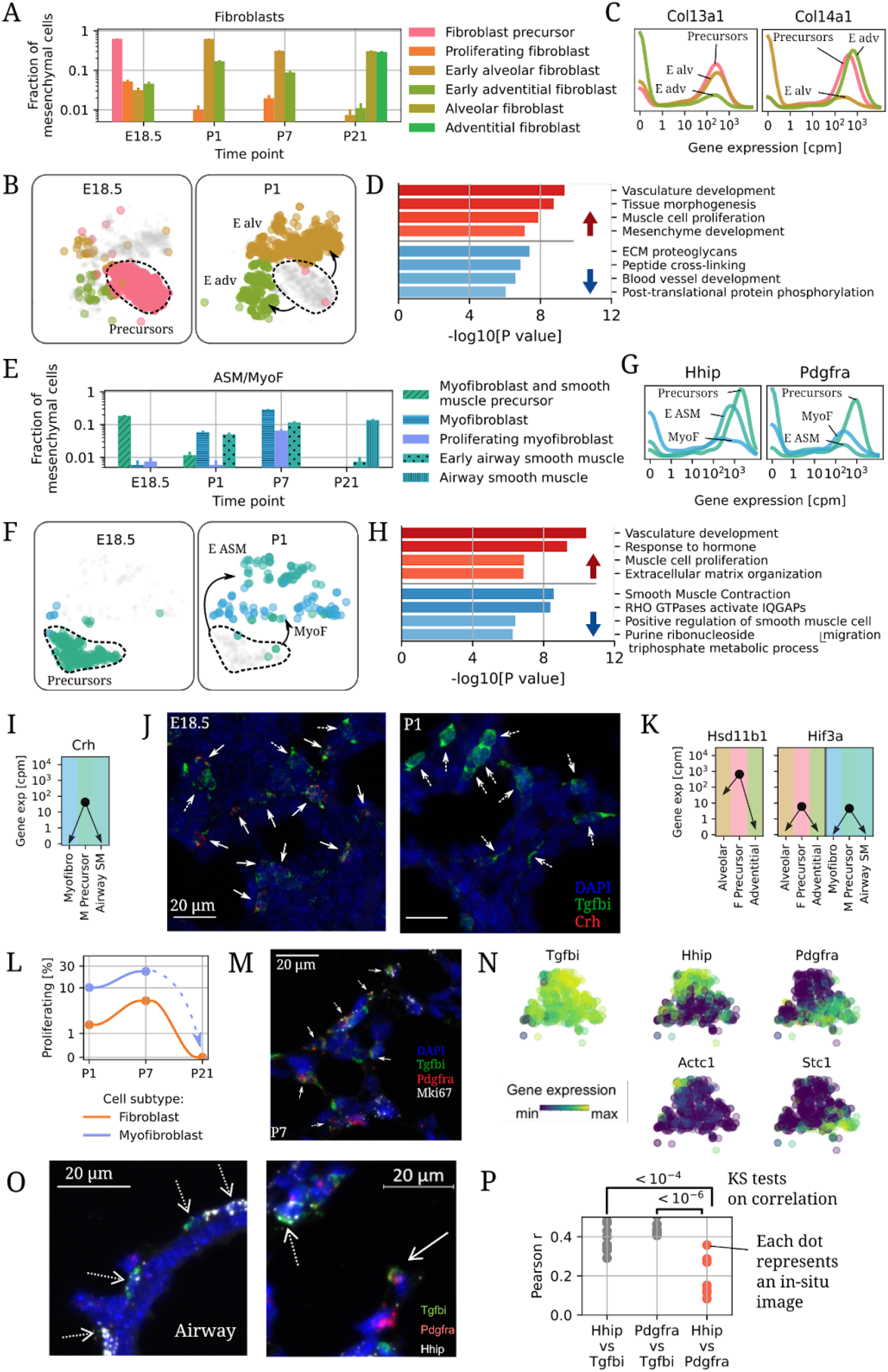
Developmental changes in perinatal fibroblasts, myofibroblasts, and airway smooth muscle. (A) Changes in composition of lung fibroblast subtypes between E18.5 and P21. (B) Embedding of fibroblasts and precursors at E18.5 (left) and P1 (right). (C) Distribution of expression for *Col13a1* and *Col14a1* in fibroblasts. (D) Pathways enriched among genes up- (red, top) or down-regulated (blue, bottom) in early postnatal fibroblasts versus prenatal fibroblast precursors [40]. (E) Changes in abundance of ASM, myofibroblasts and their precursors between E18.5 and P1. (F) Embedding of ASM and myofibroblasts at E18.5 (left) and P1 (right). (G) Distribution of expression of *Hhip* and *Pdgfra* in myofibroblasts, and early ASM, and their precursors. (H) Pathways enriched among genes up- (red, top) or down-regulated (blue, bottom) in early postnatal myofibroblasts and ASM versus their precursors. (I) Expression changes of *Crh* between E18.5 and P1. (J) In-situ hybridization of *Crh+ Tgfbi+* (solid arrows) and *Crh-Tgfbi+* cells (dashed arrows) in E18.5 and P1 lungs. (K) Expression changes of Hsd11b1 in fibroblasts and Hif3a in both fibroblasts, ASM, and myofibroblasts between E18.5 and P1. (L) Fraction of proliferative fibroblasts and myofibroblasts at postnatal time points. (M) In-situ hybridization of nonproliferating (*Mki67-*solid arrows) and proliferating (*Mki67+* dashed arrows) myofibroblasts at P7. (N) Embedding of Early ASM and myofibroblasts, colored by subtype and by expression of select genes. (O) In-situ hybridization of P7 lungs around airways (left) and in the distal lung (right). Dashed arrows: *Tgfbi+ Hhip+* cells, solid arrow: *Tgfbi+ Pdgfra+* cells. (P) Quantification of the degree of coexpression of *Hhip, Tgfbi,* and *Pdgfra* from in-situ images of P7 mice. P-values are Kolmogorov-Smirnov tests with Bonferroni multiple hypothesis correction.

ASM/MyoF precursor abundance decreased between E18.5 and P1 (**Figure 3E**) while Early ASM and myofibroblast were rare before birth and increased in abundance by P1 (**Figure 3E-F** and **Table S3**). ASM/MyoF precursors coexpressed *Hhip* and *Pdgfra,* while postnatal MyoF expressed only *Pdgfra* and postnatal Early ASM expressed only *Hhip* (**Figure 3G**). Pathway analysis suggested that ASM/MyoF precursors expressed higher levels of genes related to vascular development and muscle cell proliferation (**Figure 3H**, red bars) and lower levels of smooth-muscle contraction genes (blue bars) compared to balanced postnatal ASM/MyoF cells. ASM/MyoF precursors but not postnatal Early ASM and MyoF cells expressed corticotropin releasing hormone (*Crh*, **Figure 3I**), an early signal for glucocorticoid production within the hypothalamic–pituitary–adrenal (HPA) axis. *Crh* plays an essential role in lung maturation [41] via cortisol-induced increases in surfactant protein release [42,43]. *Crhr1* and *Crhr2*, which encode the *Crh* receptors, were not expressed by any cell type in either this or other scRNA-Seq studies on perinatal lung development (**Figure 3 Supplementary 2**) [11][18][44]. No expression of *Crh* or its receptors was detected in the adult atlas [20]. In agreement with these data, *Tgfbi+ Crh+* cells were detected by in-situ hybridization imaging (RNA-Scope) within the lung parenchyma in E18.5 mouse embryos but not postnatally (**Figure 3J**, solid arrows, n=6 animals per group). Fibroblast precursors but not postnatal fibroblasts expressed high levels of *Hsd11b1* (**Figure 3K**), the main enzyme responsible for converting cortisone into its bioactive form cortisol, indicating a local amplification of glucocorticoid action [45]. Expression of *Hif3a,* a negative regulator of *Hif1a,*was lost in both fibroblasts and ASM/MyoF cells after birth, suggesting tight regulation of this pathway at the critical transition between fetal development and air-breathing life (**Figure 3K**).

*Mki67*+ proliferating myofibroblasts and pericytes were observed at P1 and increased in abundance at P7 (**Figure 3L**). Proliferating myofibroblasts were localized throughout the parenchyma of P7 lungs as triple positive *Tgfbi*+ *Pdgfra*+ *Mki67*+ cells (**Figure 3M**, dashed arrows, n=2) adjacent to *Tgfbi+ Pdgfra+ Mki67*-nonproliferating myofibroblasts (solid arrows). There were no proliferating (myo-)fibroblasts detected via scRNA-Seq from P21 mice (**Figure 3L**), consistent with a progression towards cell quiescence towards the end of alveolarization and adulthood.

The relationship between postnatal Early ASM and myofibroblasts was then examined in detail. These cells, in addition to shared *Tgfbi* expression and exclusive expression of *Hhip* (for Early ASM) and *Pdgfra* (for myofibroblasts) as mentioned above, showed partial expression of *Actc1,* an alpha actin typically expressed in heart muscle [46], in a fraction of Early ASM and *Stc1,* a regulator of calcium levels that has been implicated in myopathy [47], in a fraction of myofibroblasts, indicating further layers of latent transcriptomic heterogeneity (**Figure 3N)**. In-situ hybridization was then used to determine the location of each cluster at P7. As expected, only *Tgfbi+ Hhip+* Pdgfra-cells, a profile matching Early ASM, were found encircling airways (**Figure 3O** left, dashed arrows, n=2), buttressing the scRNA-Seq-based annotation. Surprisingly, however, not only *Tgfbi+ Hhip-Pdgfra+* (solid arrows), matching myofibroblasts, but also *Tfgbi+ Hhip+ Pdgfra-* cells (dashed arrows), matching Early ASM, were observed in the distal parenchyma (**Figure 3O**, right, n=2). Automated image analysis on the distal parenchyma only (10 images, no airways) demonstrated a lower correlation between *Pdgfra* and *Hhip* than the correlation of either molecule with *Tgfbi* (**Figure 3P**), which mirrors the transcriptomics results (**Figure 3N**). Overall, these results support the hypothesis that while only Early ASM encircle airways at P7, both types of cells can be found in mutual proximity in the distal lung, a conclusion that could now be reached by scRNA-Seq alone.

### Mural cells change gradually during perinatal development

Mural cells, which include pericytes and vascular smooth muscle, form the outer lining of the vasculature and help support the endothelium and regulate vascular tone. In the perinatal lung, the transcriptomes of mural cells changed slowly between E18.5 and P21. Relative cell type abundance of mural cells (**Figure 1D**) and of each mural subtype (pericytes and VSM) (**Figure 4A**) increased steadily over time. A cluster of proliferating pericytes peaked in abundance at P7 (**Figure 4B**), at the same time as proliferating myofibroblasts. Confocal imaging of P7 lung confirmed the presence of proliferating pericytesthat co-expressed *Cox4i2* and *Mki67* (**Figure 4C**, n=2). A developmental shift in the pericyte transcriptome was visible in our embedding (**Figure 4D**, colors as in **Figure 1F**). Further analyses revealed specific transcriptomic progressions: genes exhibiting gradual decreases in expression over time (e.g. *Aspn),* gradual increases over time (e.g. *Enpp2),* marked downregulation after birth (e.g. *Timp3*), and either marked down-regulation (e.g. *Acta2*) or up-regulation (e.g. *Cxcl14)* between P7 and P21 (**Figure 4E**, same colors - all P-values are Bonferroni-adjusted KS tests). In contrast, VSM cells did not exhibit a clear transcriptomic shift over time in the embedding (**Figure 4F**) and lacked high expression of *Mki67* (**Figure 2A**).

**Figure 4.**
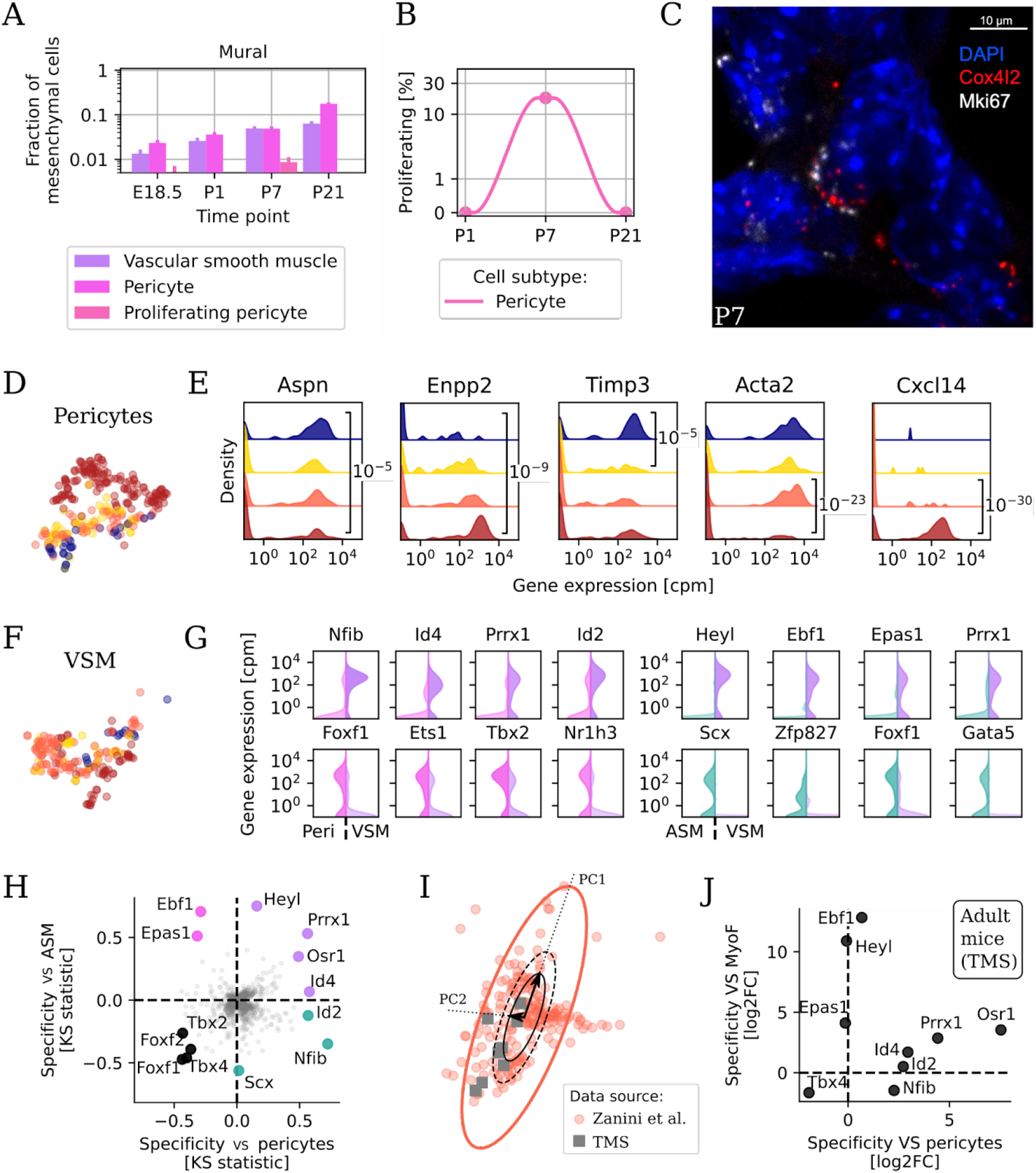
Mural cells encompass proliferating pericytes at P7 and harmonized vascular smooth muscle characterized by specific transcription factors. (A) Mural cell type composition between E18.5 and P21. Colors as in Figure 2A. (B) Fraction of proliferating pericytes at postnatal time points. (C) Representative FISH image of *Mki67+ Cox4i2+* proliferating pericytes in P7 lungs. (D) Embedding of pericytes, colored by time point. (D) Violin plots of the expression of select genes that change in expression over perinatal development in pericytes. (F) Embedding of VSM, colored by time point. (G) Violin plots of gene expression in pericytes versus VSM (left) and ASM versus VSM (right) for transcription factors (TFs) marking either cell type. (H) Combined statistical enrichment of expressed transcription factors in VSM versus pericytes (x axis) and versus ASM (y axis). Genes in the upper right quadrant (violet) are specific to VSM versus both ASM and pericytes, while genes in the lower left quadrant (black) are specifically absent in VSM but not in pericytes nor ASM. (I) Harmonized embedding of VSM from this study (red circles) and overlapping cells from Tabula Muris Senis (gray squares), identified using a rigidly expanded principle component analysis (PCA) ellipse. These cells are originally annotated as pericytes, myofibroblasts, or fibroblasts. (J) Differential expression of select transcription factors in newly identified atlas VSM versus atlas pericytes (x axis) and versus atlas myofibroblasts (y axis).

The transcriptional distinction between VSM, pericytes, and ASM remains uncertain in recent papers [19,20,28,31]. For instance, Guo et al. annotated as “Pericyte-1”a cell cluster which lacks pericyte markers (**Figure 4 Supplementary 1**). To clarify the transcriptional identity of VSM, our analysis focused on transcription factors (TFs) because they have been shown to be the best gene category to distinguish cell types [21]. Several transcription factors were upregulated in VSM (purple) relative to ASM (green) and in VSM relative to pericytes (pink) (**Figure 4G, top**) or downregulated in VSM versus the same cell types (**Figure 4G, bottom**). VSM also expressed TFs that were absent in both ASM and pericytes (**Figure 4H**, purple dots): these included *Prrx1* [48], *Osr1* [49] and *Id4* [50]. *Tbx4* [51] and other genes were expressed in both ASM and pericytes but specifically downregulated in VSM (black dots). Given the absence of a VSM annotation in adult atlas [20], we reanalyzed atlas cells that overlapped in the harmonized embedding (**Figure 2B**) with our VSM cells (**Figure 4I**). The 8 atlas cells within the principal component analysis-based ellipse (gray squares) were originally annotated as adult myofibroblast or pericyte. However, the co-embedding coordinates overlapped with those of our VSM (red circles). Differential expression of transcription factors between these 8 atlas cells and other atlas pericytes and myofibroblasts resulted in a specific TF fingerprint (**Figure 4J**) that was almost identical to the developmental VSM fingerprint previously defined (**Figure 4H**), with upregulation of *Prrx1, Osr1* and *Id4*, and downregulation of *Tbx4* compared to both other cell types. In-situ hybridization of P7 lungs with *Tgfbi, Pdgfra*, and hydrazide (which marks vessels but not airways) showed that cells lining the large airways highly expressed *Tgfbi* (which is expressed by both ASM and myofibroblasts), but cells located between the double elastic laminae of arterioles were *Tgfbi* negative (**Figure 4 Supplementary 2**). Overall, these data confirmed that VSM differ from both ASM and myofibroblasts not only in their location within the tissue but also in their transcriptional profile, in agreement with some reports [28,52].

### Hyperoxia inhibits mesenchymal transcriptional progression and pericyte proliferation

To study the discrete transcriptomic effect of hyperoxia on the developing lung, we performed scRNA-Seq and in-situ analysis on mice exposed to 80% oxygen (hyperoxia, HO) from birth through P7 [11,53]. The relative abundance of each mesenchymal cell subtype was quantified. Hyperoxia increased the relative abundance of fibroblasts but decreased the abundance of ASM, proliferating myofibroblasts and pericytes (**Figure 5A**). We hypothesized that hyperoxia might render the lung cellular composition at P7 more similar to younger animals. We compared cell type abundance profiles via bootstrapping and found that the transcriptomes of hyperoxia-exposed mice were significantly more similar to healthy P1 mice than P7 mice (P=0.001, **Figure 5B**), indicating that hyperoxia inhibited the cellular diversification of the mesenchyme. We next asked whether hyperoxia delayed developmental changes in the transcriptome within each cell type. We specifically interrogated the differential expression of genes that exhibit transcriptional progression during development in normoxia (e.g. higher at P7 than P1), and found that hyperoxia inhibited this progression in all the mesenchymal populations, especially in the mural cells (**Figure 5C**).

**Figure 5.**
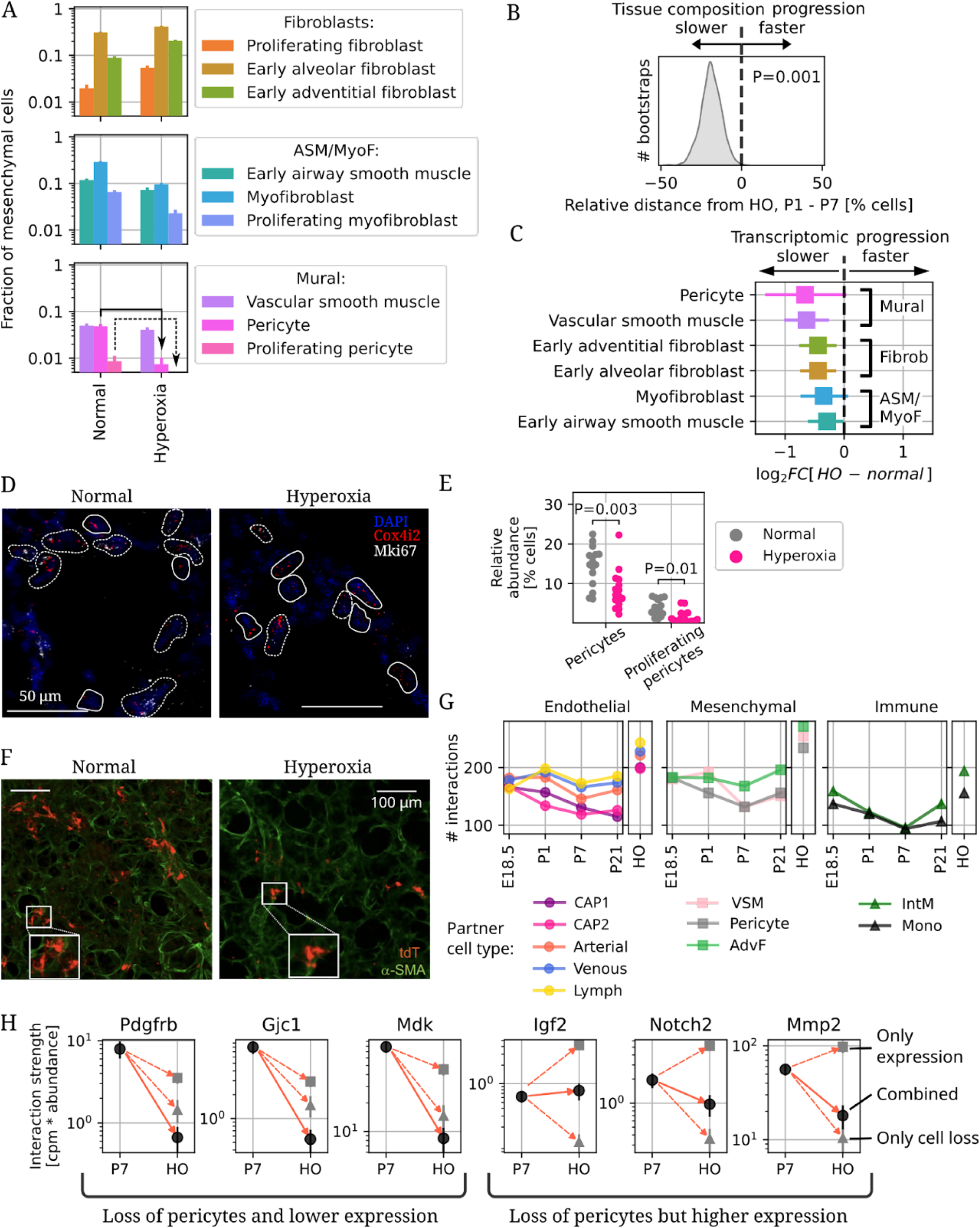
Hyperoxia delays mesenchymal transcriptional progression and suppresses pericyte proliferation. (A) Cell subtype composition in between normoxia and hyperoxia at P7. (B) Cell subtype composition changes in hyperoxic mice. The arrows indicate the observed severe loss of pericytes (solid) and complete loss of proliferating pericytes (dashed). (B) Kernel density estimate of 1,000 bootstraps over cells (balanced at each time point) for the difference in distance on cell composition between hyperoxic mice and P7, minus hyperoxic mice and P1. A positive number indicated that the hyperoxic mesenchyme composition is closer to a healthy P1 mouse than a P7 one. Only 1 out of 1000 simulations showed hyperoxia closer to P7 than P1. (C) Differential expression between normoxic and hyperoxic cells in each cell type for genes progressing during early postnatal development (upregulated at P7 versus P1). (D) Representative in-situ images of pericytes (solid) and proliferating pericytes (dashed) in P7 normoxia and hyperoxia mice. (E) Quantification of the relative abundance of pericytes and proliferating pericytes in in-situ images from P7 mice using an unsupervised image analysis approach over thousands of cells. (F) Representative confocal imaging of lung tissue at P7 from Notch3-CreERT2-tdT expressing mouse pups who received tamoxifen from P1-P3 and were placed in either normoxia or hyperoxia for seven days from birth to P7. α-smooth muscle actin (α-SMA) shown in green. (G) Number of detected ligand-receptor cell-cell interactions between pericytes and other cell types at each time point during normal development and in hyperoxia mice at P7. (H) Strength of select interactions with pericytes measured as product of gene expression and pericyte abundance in normoxia and hyperoxia. Data sources: endothelial cells [18] and immune cells [11].

Remarkably, the scRNA-Seq data indicated that pericytes and especially proliferating pericytes were almost completely lost in hyperoxia mice (**Figure 5A**, solid and dashed arrows, respectively). This marked decrease in pericytes and proliferating pericytes was confirmed *in situ* by detecting the combination of *Mki67* and *Cox4i2* in hyperoxia-exposed lungs (**Figure 5D**, n=2 for each group). Quantification of the results using automated image segmentation and analysis demonstrated that hyperoxia reduced the abundance of both total pericytes (P=0.003, KS test) and proliferating pericytes (P=0.01, KS test) (**Figure 5E**, each dot represents an analyzed image). To further confirm this observation Notch3-CreERT2-tdT mice were given tamoxifen at P0 and P3 and placed in normoxia or hyperoxia from until P7. Deep imaging of thick distal lung sections (150 μm) indicated that compared to normoxia, pericyte number was markedly reduced in hyperoxia (**Figure 5F**, representative images of n=3 animals for each group), and the morphology of the pericytes disrupted, with a decrease in the number and length of projections (**Figure 5F**, high mag inset).

We then asked whether the loss of pericytes might impact the cell-cell communication networks used by these cells to help orchestrate postnatal lung development. We reused the CellPhoneDB database [54] to count the number of putative ligand-receptor interactions (at least 20% of cells expressing each gene) between pericytes and EC, other mesenchymal cells, and immune cells at all four time points and in hyperoxia mice at P7. Across all partner cell types, hyperoxia increased the number of putative interactions with pericytes (**Figure 5G**), suggesting that the organism might be compensating for the partial loss of pericytes with a global transcriptional upregulation of communication-related genes in both pericytes and the partner cell types. To examine this hypothesis in closer detail, we quantified for select genes their overall “interaction strength” in pericytes, i.e. the product of pericyte abundance times gene expression level in normoxia and hyperoxia (**Figure 5H**). For some genes, down-regulation intensified the effect of pericyte loss, possibly leading to loss of signaling. This phenomenon was observed for *Pdgfrb*, an essential component of a pathway promoting pericyte proliferation and migration towards the endothelium during vascular development [55], *Gjc1,* a connexin that promotes vessel stability and EC quiescence [56], and midkine or *Mdk,* a heparin binding, proangiogenic cytokine [57]. In contrast, other genes exhibited up-regulation of gene expression, which may serve to mitigate the effect of diminished pericyte type abundance and thereby preserve signaling.

### Hyperoxia can cause the emergence of *Acta1*+ cells in the neonatal lung

In one (male) hyperoxia mouse, novel populations of *Acta1*+ cells were observed (**Figure 6A**), which had not been detected in normal development by us or other studies and had not been described in a recent study on hyperoxia [19]. The two novel clusters (henceforth called HA1 and HA2) were projected in distinct parts of the embedding, with HA2 located close to Alveolar fibroblasts (**Figure 6B**). Both clusters expressed the neuron-specific tubulin (*Tubb3*) and *Lgals3*, encoding a -galactosidase binding lectin that is up-regulated in tissue fibrosis [58] as well as the cardiac muscle gene *Actc1* and the membrane channel *Aqp3*, and one cluster (HA2) also expressed genes characteristic of alveolar fibroblasts (e.g. *Wnt2, Col13a1*) (**Figure 6C**), explaining its embedding proximity to fibroblasts. *Acta1+* cells lacked expression of genes marking endothelial (e.g. *Pecam1, Cdh5*), epithelial (e.g. *Epcam*), or immune cells (e.g. *Ptprc*).

**Figure 6.**
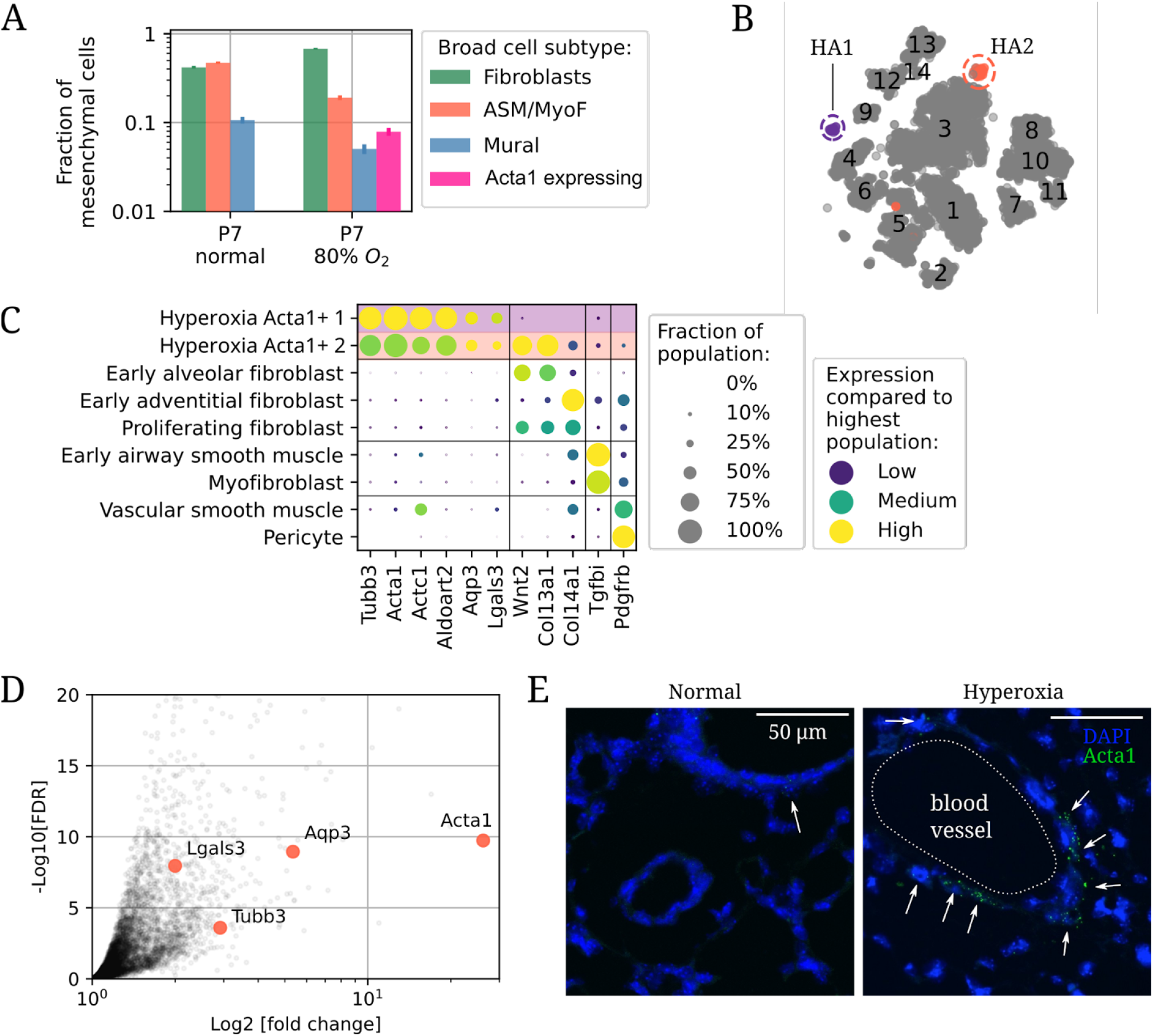
Hyperoxia causes the emergence of *Acta1+* cells in the lung. (A) Relative cell subtype abundance for broad mesenchymal types in normoxia and hyperoxia. (B) Embedding as in **Figure 2A** but with the additional populations hyperoxic *Acta1+* 1 (HA1, in purple) and hyperoxic *Acta1+* 2 (HA2, in red). Other cells are colored in gray. (C) Dot plot of marker genes for *Acta1+* cells. (D) Reanalysis of bulk transcriptomics in P7 lungs exposed to hyperoxia shows up-regulation of *Acta1* and other marker genes for the novel cell populations. Data from [59]. FDR: False discovery rate. Downregulated genes are not shown. (E) Representative *in-situ* hybridization images of *Acta1* (green) expression in normoxia and hyperoxia. Cells with three or more *Acta1* molecules are indicated by arrows.

Given *Acta1+* cells were initially observed in a single animal, we sought to validate their presence via additional experiments on multiple, distinct mice. Reanalysis of bulk RNA-Seq data from an independent study on hyperoxia-exposed P10 murine lungs confirmed an upregulation of *Tubb3, Acta1, Aqp3,* and *Lgals3* compared to normal P10 mice [59] (**Figure 6D**). Moreover, *In-situ* hybridization imaging of lungs from a separate group of mice (n=4 for each group) confirmed an increased density of *Acta1+* cells in both male and female hyperoxia-exposed mice versus normal P7 mice, especially near pulmonary blood vessels (**Figure 6E**). Taken together, these results validate the proposition that hyperoxia can be but is not always accompanied by the appearance of previously undetected but transcriptionally distinct *Acta1+* cells in the neonatal lung.

## Discussion

Mesenchymal cells are responsible for essential functions in the developing lung, including modulating the composition of the extracellular matrix, providing contractility to airways and vasculature, and supporting angiogenesis and alveolarization [5,8,28]. Single-cell transcriptomics has been recently applied by several groups to investigate the composition of the lung mesenchyme with high granularity [23,28,31,52]. Despite similar experimental approaches across studies, the computational analyses and resulting biological interpretations have differed greatly, creating an uncertain landscape in terms of cell type annotation and transcriptional changes across development. This study aimed to establish a coherent biologic description of the perinatal lung mesenchyme that could overcome the discrepancies in the existing data on normal development, and subsequently use it as a foundation to investigate the effect of hyperoxia on the cellular composition of the lung and gene expression within each mesenchymal cell subtype.

The transcriptional identity of several mesenchymal cell subtypes was clarified. Vascular smooth muscle cells could be assigned a distinct transcriptomic signature from airway smooth muscle, myofibroblasts, and pericytes, with which they have been previously confused [20,21,23,31]. Two types of fibroblasts, alveolar and adventitial, but no lipofibroblasts were seen, as clearly demarcated by some [21] but not all papers [28,31,52]. Myofibroblasts and airway smooth muscle cells could be attributed distinct transcriptional profiles during early postnatal development but only airways smooth muscle cells were detected later at P21, questioning a recent hypothesis that a population of ductal myofibroblasts persists through adulthood [52]. Intriguingly, the identification of an embryonic mesenchymal cell population expressing *Crh,* which encodes a hormone that controls systemic glucocorticoid levels via the hypothalamus - pituitary gland - adrenal gland axis, was identified computationally and *in situ,* clarifying brief mentions in previous literature [28,52]. Mouse embryos from *Crh*-deficient mothers fail to generate sufficient airspaces around E17.5 to E18.5 while newborn pups die within 24 hours [60]. In the light of the present results, these physiologic deficiencies might stem from failure to produce and likely secrete *Crh* by myofibroblast precursors in the lung itself, suggesting these cells could act as a master regulator of airspace formation during late embryonic development. Future studies employing cell type-specific *Crh* knockout within these embryonic precursors will be key to test this hypothesis. Moreover, additional research is needed to determine whether the pituitary or adrenal glands, or even other tissues, are involved in the *Crh*-driven regulation of glucocorticoid levels in the neonatal lung.

While the effects of hyperoxia on the transcriptomes of lung cells have been described in previous publications [18,19], this study focused specifically on the effects on hyperoxia on mesenchymal cells. The dramatic loss of proliferating pericytes and the subsequent alteration of cell-cell communication networks are suggestive of a potentially underappreciated role for pulmonary pericytes in protecting against neonatal lung injury [61]. While the specific functions of lung pericytes are not fully understood, pericyte deficiency can disrupt the blood-brain barrier and contributes to the capillary instability, vascular leak and macular edema of diabetic retinopathy [62,63]. Ongoing efforts by our groups are aimed at clarifying the physiologic implications of partial pericyte loss on the altered tissue structure and pathologic vascular remodeling observed during lung recovery after hyperoxia. Given that this murine model recapitulates many features of the human disease, elucidation of novel functions for understudied cell types might lead to new strategies to improve clinical care for prematurely born infants with bronchopulmonary dysplasia.

The emergence of *Acta1*-expressing cells in multiple hyperoxia-exposed mice, seen by single cell and bulk transcriptomics and confirmed via in-situ hybridization, raises a number of questions about their developmental ontogeny as well as their pathophysiologic function. While much remains to be discovered about these cells, in a previous study cardiomyocyte-associated genes were differentially expressed in a bulk transcriptomic of whole lung after hyperoxia exposure [64]. Intriguingly, cardiomyocytes of the pulmonary vein, which have been associated with asthma [65], express *Actc1* and partially *Acta1* but not *Tubb3* and *Aqp3* [24], suggesting that *Acta1*-expressing cells might be functionally related and possibly developmentally linked to cardiomyocytes. Given the large variability in severity seen in both hyperoxia-exposed mice and infants with bronchopulmonary dysplasia - only one of our two mice sequenced via single cell RNA-Seq contained *Acta1+* cells -, it is tempting to speculate that these cells might be a stochastic response of the body restricted to the most severe forms of disease. Future experiments using transgenic mice for *Acta1, Tubb3,* or other marker genes specific for these cells are planned to test this hypothesis.

Overall, this study constitutes a coherent cellular portrait of the perinatal lung mesenchyme that aims to overcome discrepant biologic interpretations in the existing literature and deeply characterize the cellular and transcriptional changes that accompany neonatal hyperoxia exposure, a widely used model for bronchopulmonary dysplasia. The combination of *de novo* high-quality data collection, integration with and comparative secondary analysis of extant data sets, and validation via in-situ hybridization provides a blueprint that could be used to systematically improve our understanding of the mesenchymal compartment across organs and organisms.

## Methods and Materials

### Reagents table

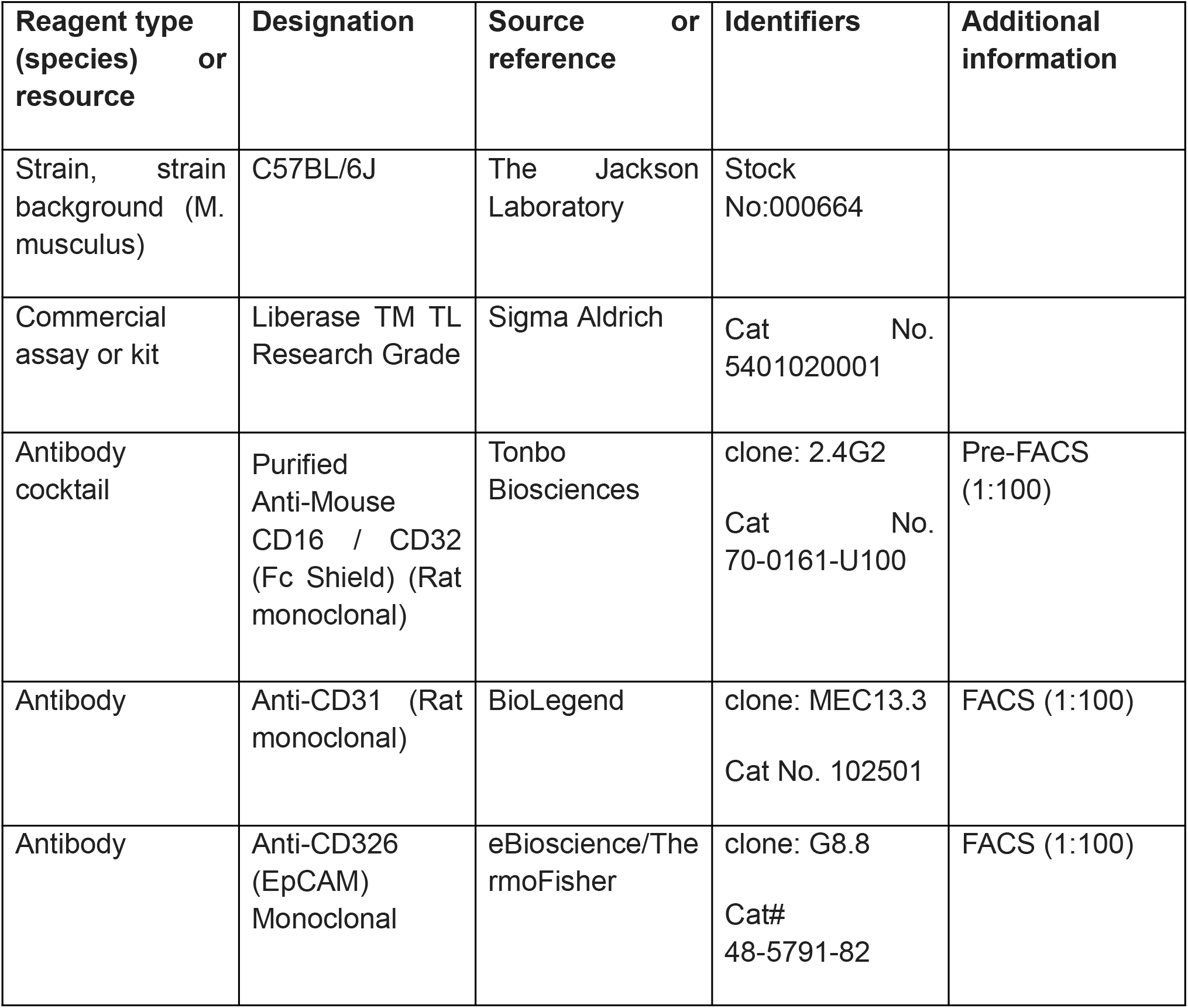
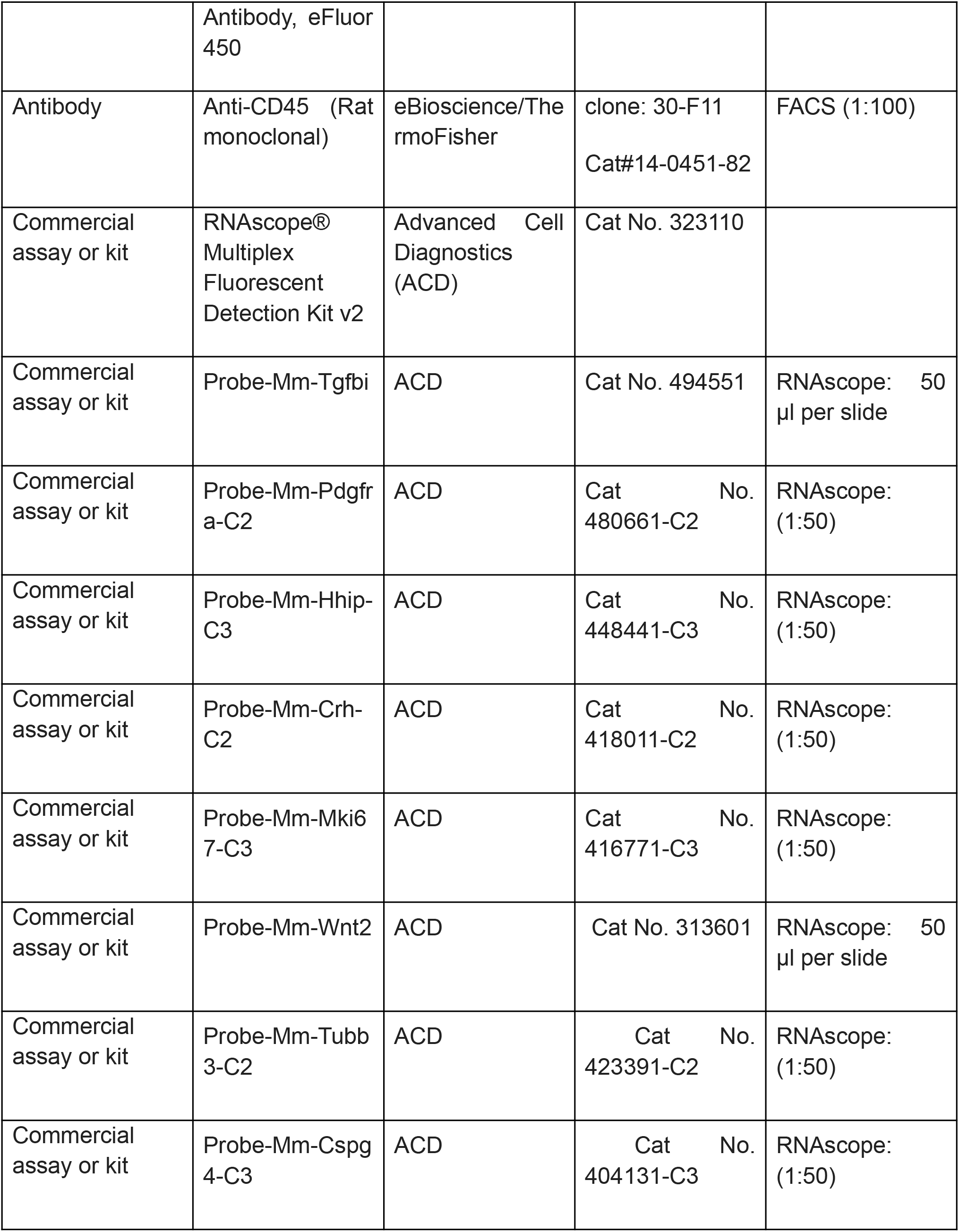
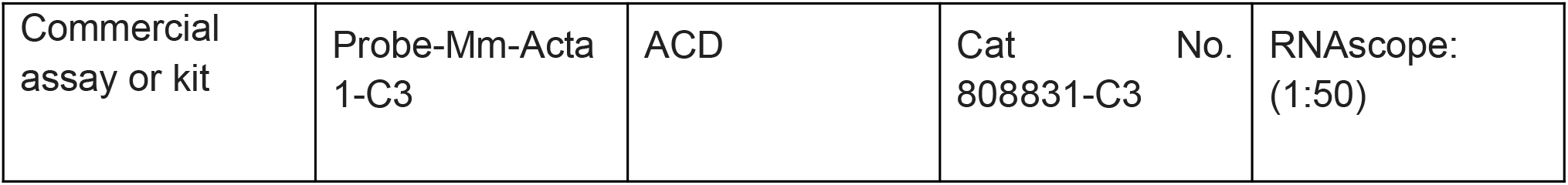

### Mouse lung cell isolation

C57BL/6 mice were obtained from Charles River Laboratories. For studies using E18.5, P1, and P7 murine lungs, pregnant dams were purchased, and pups aged prior to lung isolation. At E18.5, the dam was asphyxiated with CO2 and pups extracted. At P1, P7, and P21 pups were euthanized with euthanasia solution (Vedco Inc). For hyperoxia experiments, pups were exposed to 80% atmosphere for 7 days as previously described [18]. Genetic sex of mice at developmental stages E18.5, P1 and P7 was determined by performing PCR amplification of the Y chromosome gene Sry. P1 and P7 mice were sexed through identification of a pigment spot on the scrotum of male mice [66]. For all timepoints, female and male mice were randomly selected for the studies. For all timepoints, except E18.5, the pulmonary circulation was perfused with ice cold heparin in 1x PBS until the circulation was cleared of blood. Lungs were minced and digested with Liberase (Sigma Aldrich) in RPMI for 15 (E18.5, P1, and P7) or 30 (P21) minutes at 37 C, 200 rpm. Lungs were manually triturated and 5% fetal bovine serum (FBS) in 1x PBS was used to quench liberase solution. Red blood cells were lysed with 1x RBC lysis buffer (Invitrogen) as indicated by the manufacturer and total lung cells counted on Biorad cell counter (BioRad). Protocols for the murine studies adhered to American Physiological Society/US National Institutes of Health guidelines for humane use of animals for research and were prospectively approved by the Institutional Animal Care and Use Committee at Stanford (APLAC #19087).

### Immunostaining and fluorescence-activated cell sorting (FACS) of single cells

Lungs were plated at 1 × 10^6^ cells per well and stained with Fc block (CD16/32, 1:100, Tonbo Biosciences) for 30 min on ice. Cells were surface stained with the endothelial marker CD31 (1:100, eBioscience/ThermoFisher), epithelial marker Epcam (1:100, eBioscience/ThermoFisher), and immune marker CD45 (1:100, eBioscience/ThermoFisher) for 30 min on ice. The live/dead dye, Sytox Blue (Invitrogen), was added to cells and incubated for 3 min prior to sorting into 384-well plates (Bio-Rad Laboratories, Inc) prefilled with lysis buffer using the Sony LE-SH800 cell sorter (Sony Biotechnology Inc), a 100 μm sorting chip (Catalog number: LE-C3110) and ultra-purity mode. Single color controls were used to perform fluorescence compensation and generate sorting gates. 384-well plates containing single cells were spun down, immediately placed on dry ice and stored at −80°C.

### cDNA library generation

RNA from sorted cells was reverse transcribed and amplified using an adaptation of the Smart-Seq2 protocol for 384-well plates [11]. Concentration of cDNA was quantified using Quant-it Picogreen (Life Technologies/Thermo Fisher) to ensure adequate cDNA amplification and cDNA was normalized to 0.4 ng/uL. Tagmentation and barcoding of cDNA was prepared using in-house Tn5 transposase and custom, double barcoded indices [21]. Library fragment concentration and purity were quantified by Agilent bioanalyzer. Libraries were pooled and sequenced on Illumina NovaSeq 6000 with 2 × 100 base kits and at a depth of around 1 million read pairs per cell.

### Data analysis and availability

Sequencing reads were mapped against the mouse genome (GRCm38) using <u>STAR aligner</u> [67] and gene expression was quantified using <u>HTSeq</u> [68]. To coordinate mapping and counting, <u>snakemake</u> was used [69]. Gene expression count tables were converted into loom objects (https://linnarssonlab.org/loompy/) and cells with less than 50,000 uniquely mapped reads or less than 400 genes per cell were discarded. Doublets were discarded by excluding small clusters and single cells that coexpress markers for incompatible cell types. Counts for the remaining NNN cells were normalized to counts per million reads. For t-distributed stochastic embedding (t-SNE) [13], 500 features were selected that had a high Fano factor in most mice, and the restricted count matrix was log-transformed with a pseudocount of 0.1 and projected onto the top 25 principal components using <u>scikit-learn</u> [70]. Unsupervised clustering was performed using <u>Leiden</u> (C++/Python implementation) [25]. Singlet (https://github.com/iosonofabio/singlet) and custom Python3 scripts were used: the latter are available at https://github.com/iosonofabio/lung_neonatal_mesenchymal/. Pathway analysis on the differentially expressed genes was performed via Metascape [40] on the top 100 most differentially expressed genes for each comparison: precursors versus sample-balanced joint progenies and subsequent time points within each cluster. The most enriched pathways against a permutation test are shown ordered by significance from top to bottom (negative log of the P-value). Batch-corrected KNN [26] was used to compare our P21 data with the Smart-seq 2 data from Tabula Muris Senis [20]. Raw fastq files, count tables, and metadata are available on NCBI’s Gene Expression Omnibus (GEO) website: GSE172251 and GSE175842 (immune cells). An h5ad file with the processed data is available on FigShare at https://figshare.com/articles/dataset/Cell_atlas_of_the_murine_perinatal_lung/14703792. Bulk transcriptomic data were sourced from and are available in [59].

### In-situ validation using RNAscope and immunofluorescence (IF)

Embryonic and post-natal mice were euthanized as described above. Female and male mice were randomly selected from the litter, and at least two litters were used to source the lung tissue for all validation studies. E18.5 lungs were immediately placed in 10% neutral buffered formalin following dissection. P1, P7, and P21 murine lungs were perfused as described above, and P7 and P21 lungs inflated with 2% low melting agarose (LMT) in 1xPBS and placed in 10% neutral buffered formalin. Following 20 hr incubation at 4C, fixed lungs were washed twice in 1xPBS and placed in 70% ethanol for paraffin-embedding. In situ validation of genes identified by single cell RNA-seq was performed using the RNAscope Multiplex Fluorescent v2 Assay kit (Advanced Cell Diagnostics) and according to the manufacturer’s protocol. Formalin-fixed paraffin-embedded (FFPE) lung sections (5 μm) were used within a day of sectioning for optimal results. Nuclei were counterstained with DAPI (Life Technology Corp.) and extracellular matrix proteins stained with hydrazide [71]. Opal dyes (Akoya Biosciences) were used for signal amplification as directed by the manufacturer. Images were captured with Zeiss LSM 780 and Zeiss LSM 880 confocal microscopes, using 405 nm, 488 nm, 560 nm and 633 nm excitation lasers. For scanning tissue, each image frame was set as 1024 × 1024 and pinhole 1AiryUnit (AU). For providing Z-stack confocal images, the Z-stack panel was used to set z-boundary and optimal intervals, and images with maximum intensity were processed by merging Z-stacks images. For all both merged signals and split channels were collected.

### Image quantification

Images were quantified in two ways. First, for pericytes we used automatic image segmentation using cellprofiler [72] to identify RNA molecules from RNA-Scope and the cell boundaries, and then counted the number of cells with at least 3 positive RNA molecules in different conditions. Second, for other cells that were challenging to segment (e.g. ASM/MyoF), whole-image correlations of the fluorescence channels were computed.

### In-situ validation of pericyte abundance using transgenic mice

Notch3-CreERT2-tdT mice were given tamoxifen at P0 and P3 and placed in normoxia or hyperoxia (0.80 FiO2) from P1 to P7. Lungs were harvested at P7 and inflation fixed. Deep imaging of thick lung sections (150 μm) allowed us to visualize the abundance and location of tdT-labeled pericytes in the distal lung.

### Statistical analyses

Single cell omics data do not follow a normal distribution and are often difficult to parameterize altogether. Therefore, to ensure statistical soundness of all analyses, a statistical plan based on nonparametric tests was deployed throughout the study, exploiting the fact that hundreds of quasi-replicate cells were sampled. To identify differentially expressed genes within cell populations, between time points, and between normoxia and hyperoxia, nonparametric Kolmogorov-Smirnov (KS) tests on the distributions of gene expression were performed, and either the genes with the largest absolute value of the test statistic or the genes above a certain KS statistic (e.g. 0.3) with the largest average fold change were chosen. Nonparametric bootstrapping was used to evaluate the developmental arrest of hyperoxia. Correlation tests and proportion tests for cell population abundances were used in some image analyses.

## Acknowledgements

We thank Sai Saroja Kolluru (Stanford University) for assistance with library submission to the Chan Zuckerberg Biohub, Yuan Xue (Stanford University) for assistance with the initial single cell RNA-seq data acquisition, Astrid Gillich for technical support with the RNAscope experiments, and Maya Kumar for providing hydrazide. We also thank the Stanford Shared FACS Facility, Lisa Nichols, Meredith Weglarz, and Tim Knaak for assistance with the flow cytometry instrumentation and antibody panel design. Flow cytometry data was collected on an instrument in the Stanford Shared FACS Facility obtained using NIH S10 Shared Instrument Grant (S10RR027431-01). This work was supported by National Institutes of Health grants HL122918 (CMA), HL160018, HD092316 (CMA, DNC), the Stanford Maternal Child Health Institute Tashia and John Morgridge Faculty Scholar Award (CMA), the Stanford Center of Excellence in Pulmonary Biology (DNC), Bill and Melinda Gates Foundation (SRQ), and the Chan Zuckerberg Biohub (DNC. and SRQ). MAS is supported by the NSF-GRFP. NES was supported by a Stanford Maternal and Child Health Research Institute Ernest and Amelia Gallo Endowed Postdoctoral Fellowship and Chan Zuckerberg Biohub Physician-Scientist Fellowship.

## Notes

### Competing Interest Statement

The authors have declared no competing interest.

### Summary of Updates

We have edited the manuscript to improve readability, adapted figures, and performed additional analyses and experiments.

