## Supplementary Figures for "Lung mesenchymal cell diversity rapidly increases at birth and is profoundly altered by hyperoxia"

Supplementary Materials (for each figure)

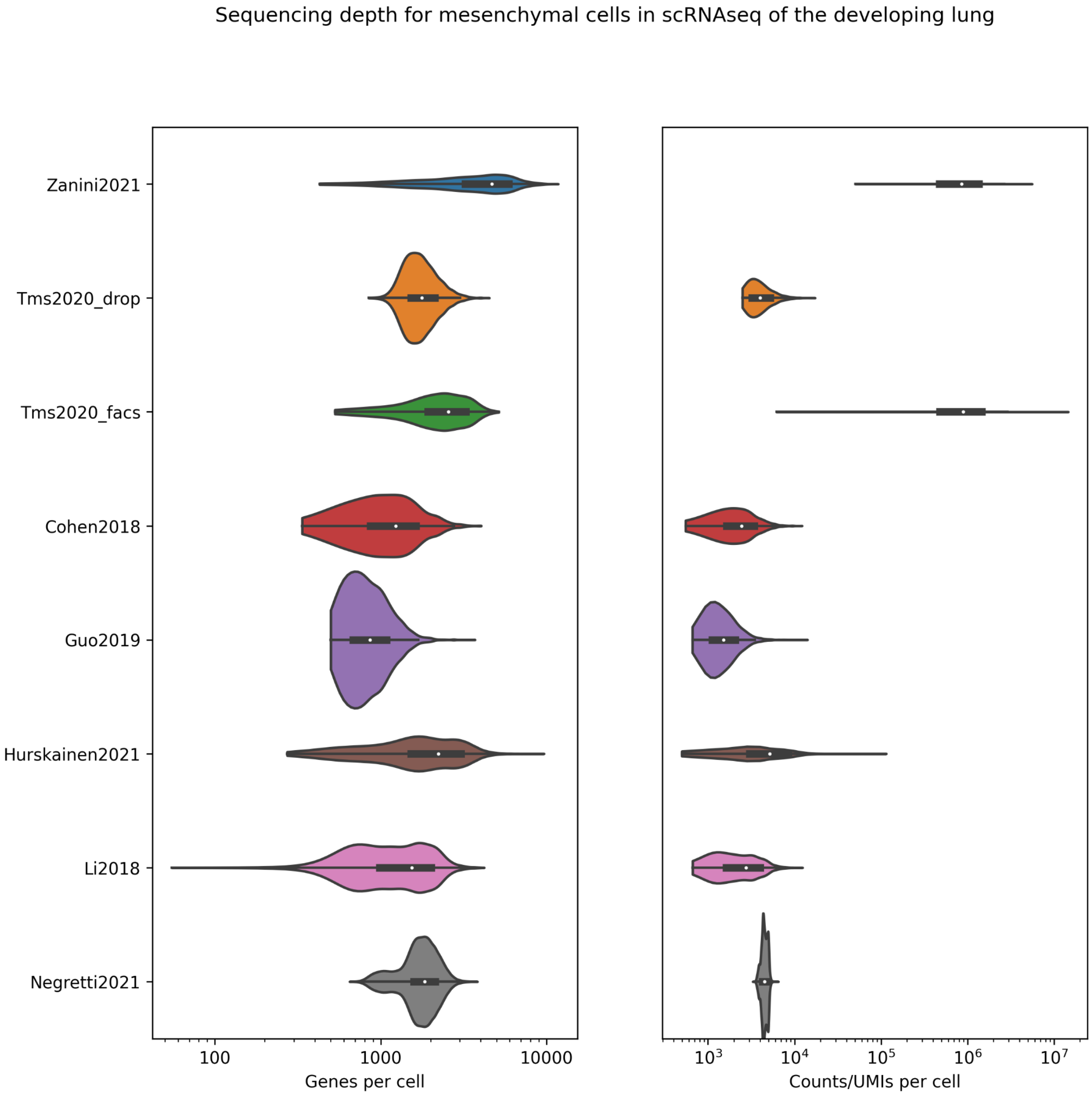


**Figure 1 Supplementary 1. Violinplot with numbers of genes per cell and number of reads/UMIs per cell for mesenchymal cells from several scRNA-Seq data sets from murine lungs.**

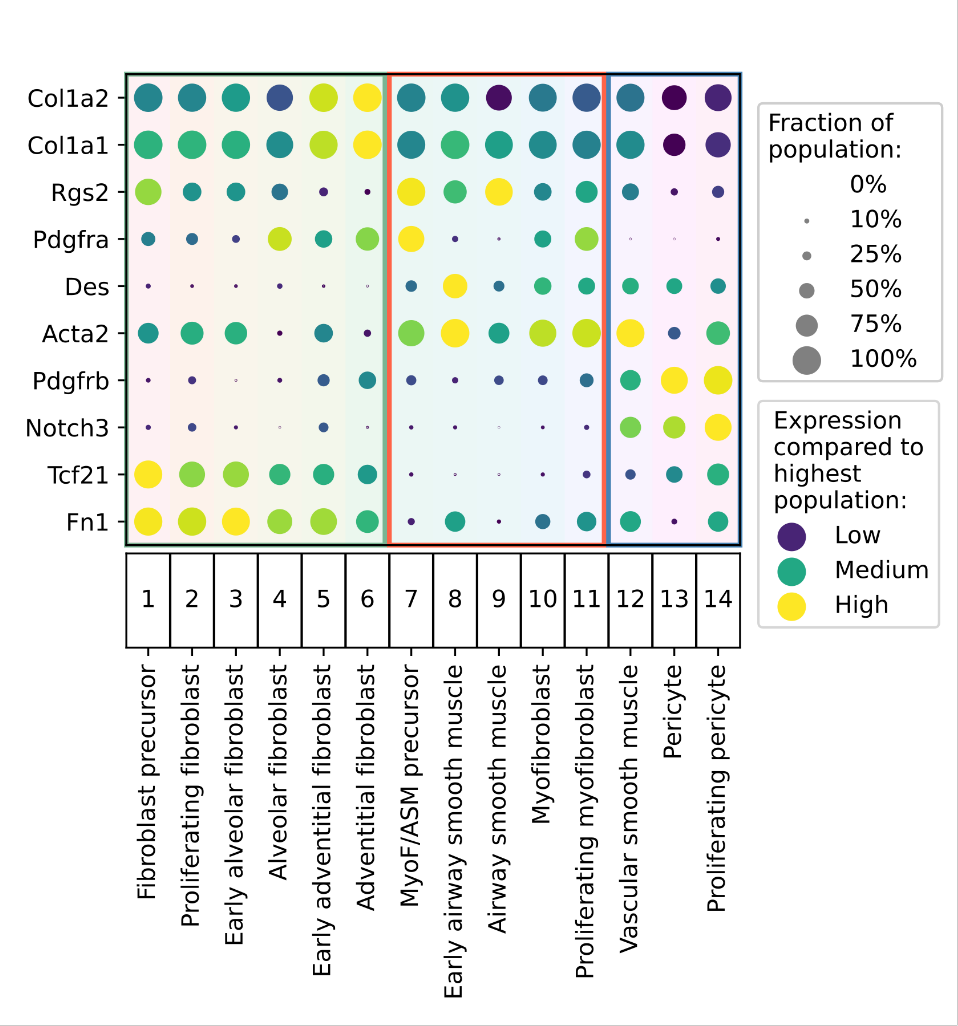


**Figure 2 Supplementary 1. Dot plot of the marker genes by Guo et al. across our subtypes.** These genes are the same as in ref [31], Figure 1b, limited to the markers for their mesenchymal populations.

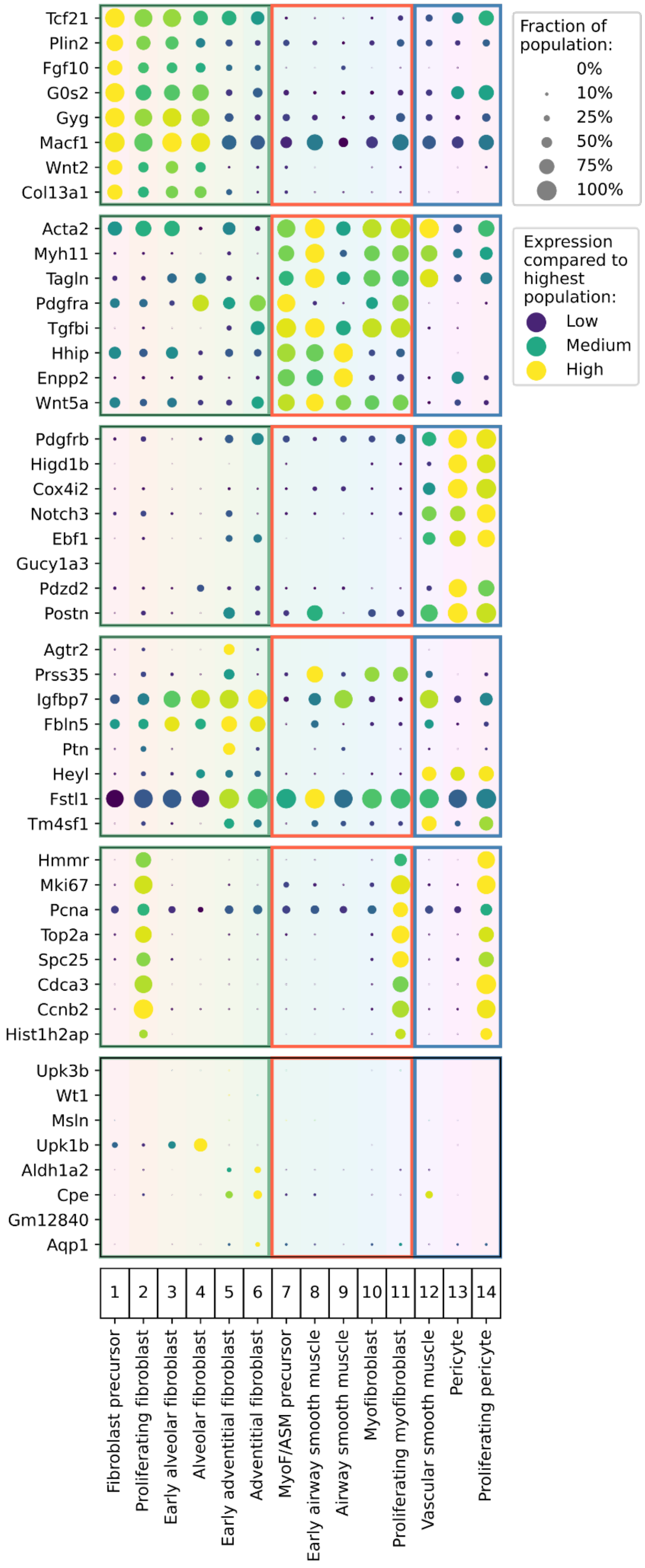


**Figure 2 Supplementary 2. Dot plot of the marker genes by Liu et al. across our subtypes.** These genes are the same as in ref [23], Figure 1E.

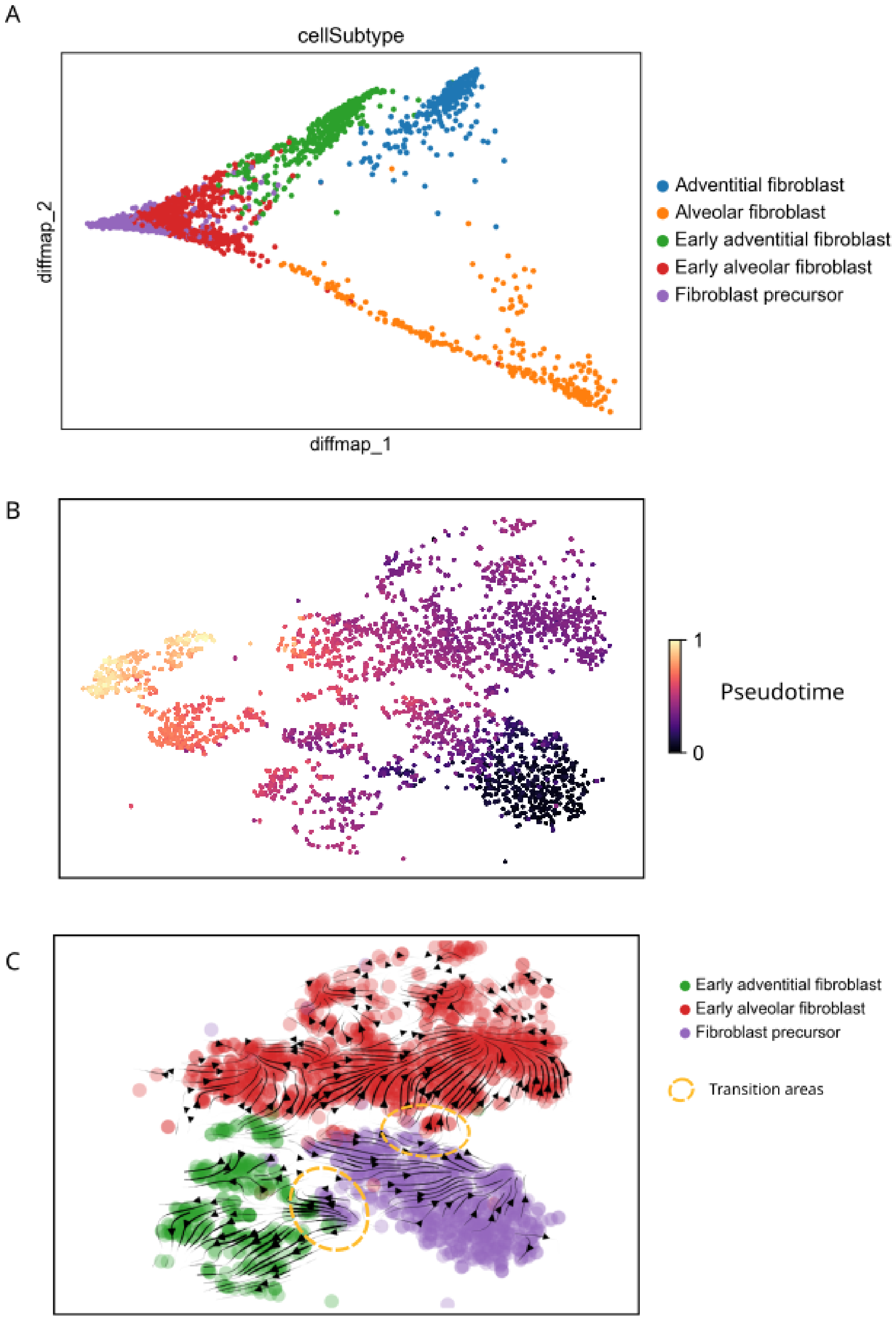


**Figure 3 Supplementary 1. Trajectory analysis of fibroblast precursors and early postnatal fibroblasts.** (A) Diffusion map colored by MS subtype. (B) Pseudotime analysis (Wolf et al., Genome Biology 2019) starting from fibroblast precursors. (C) RNA velocity stream analysis using scVelo Bergen *et al.* Nature Biotech 2020) with transitional areas between precursors and postnatal fibroblasts highlighted.

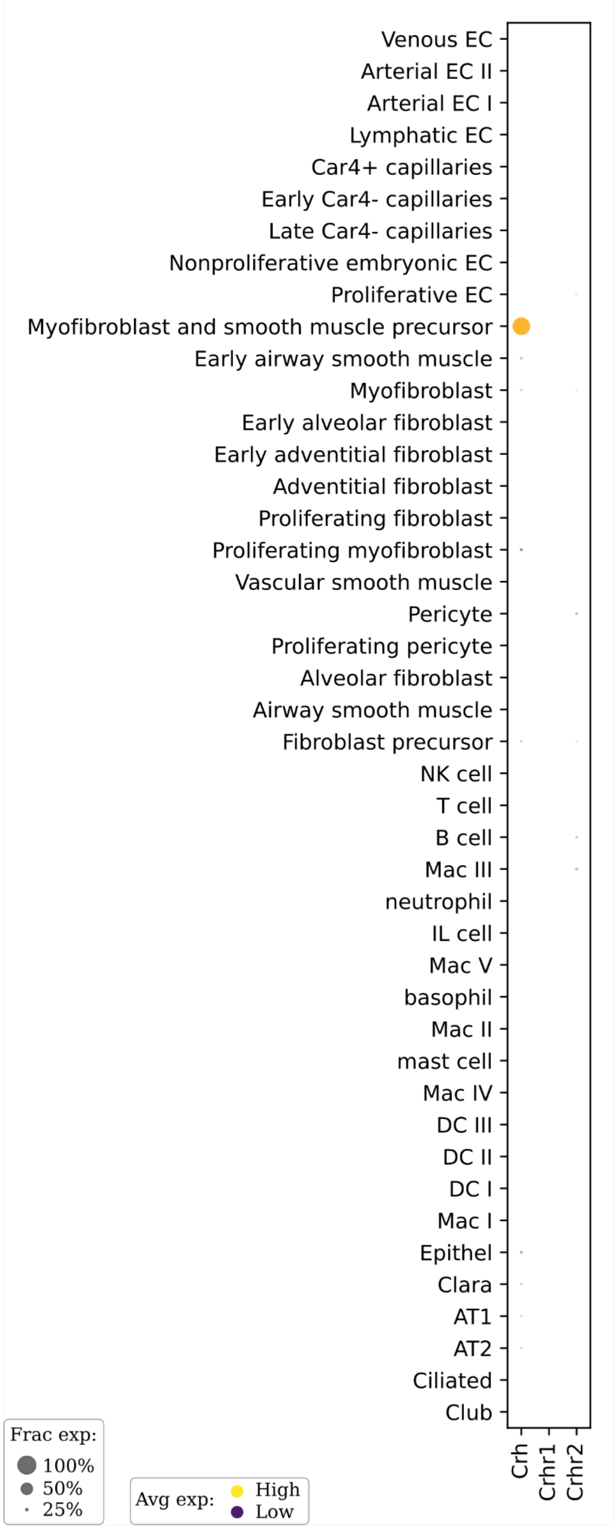


**Figure 3 Supplementary 2. Dot plot of Crh and its receptors in developing murine lungs.** Mesenchymal cell data from this article, endothelial cells from [18], immune cells from [11], epithelial from [44]. Neither of the receptors is expressed in the developing lung in any detected cell type.

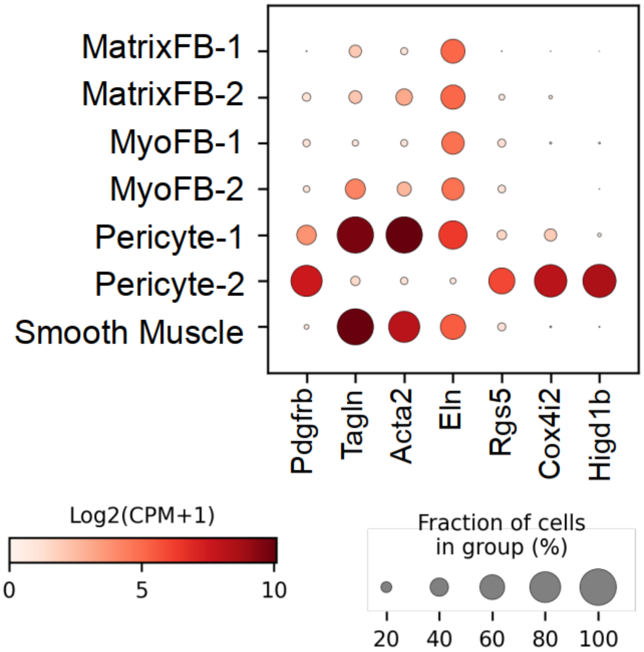


**Figure 4 Supplementary 1: Dot plot for pericyte and VSM marker genes in Guo et al [31].**

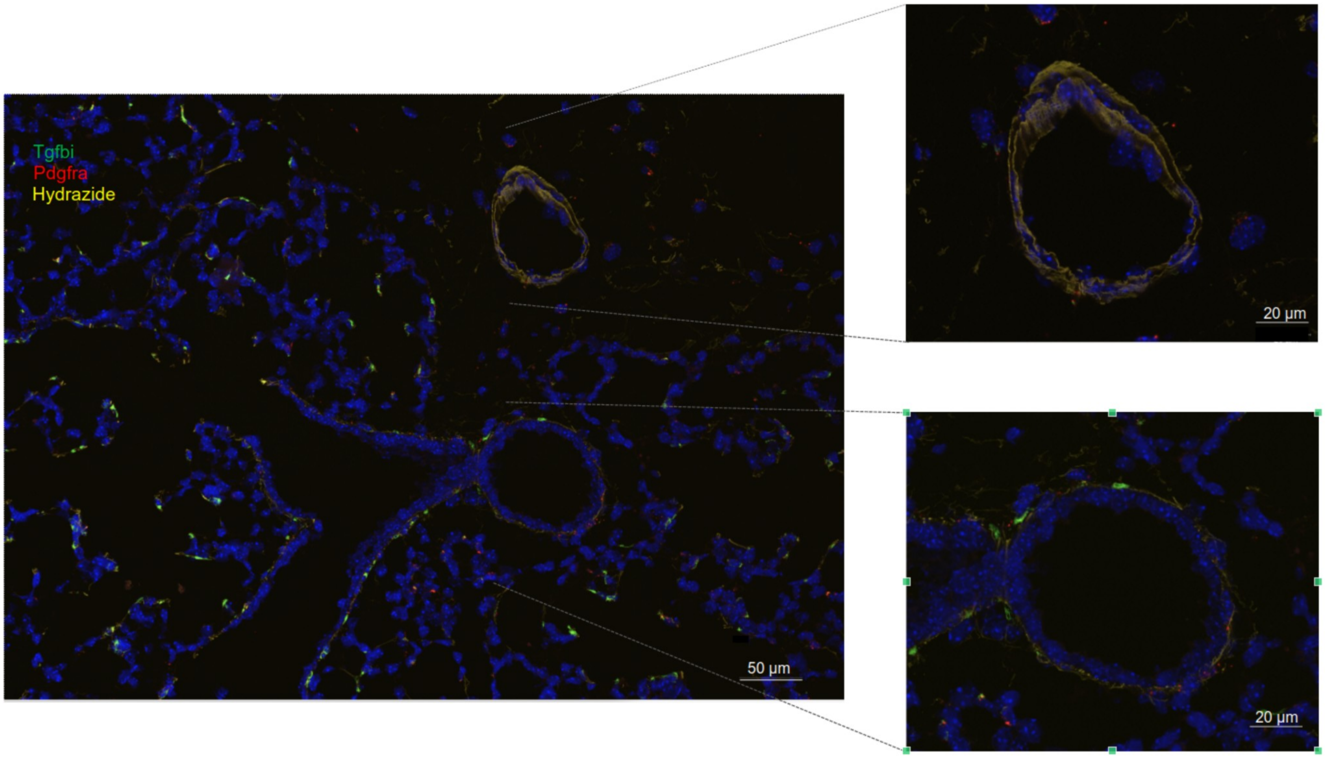

**Figure 4 Supplementary 2**. *In situ* hybridization of lung at P7 to detect Tgfbi (green), a marker for ASM/MyoF but not VSM, Pdgfra (red) and hydrazide (yellow). The enlarged zoom-ins show a blood vessel (top) and an airway (bottom).
